# Treatment Failure among People living with HIV taking Antiretroviral Therapy in Ethiopia

**DOI:** 10.1101/577049

**Authors:** Yimam Getaneh, Atsbeha G Egziabhier, Kidist Zealiyas, Rahel Tilahun, Mulu Girma, Gebremedihin G Michael, Tekalign Deressa, Ebba Abate, Desta Kassa, Yibeltal Assefa

**Affiliations:** Ethiopian Public Health Institute, HIV/AIDS and TB Research Directorate, Addis Ababa, Ethiopia; School of public health, the University of Queensland, Australia

**Keywords:** Antiretroviral therapy, treatment failure, virologic failure, viral load suppression

## Abstract

**Background:** Although treatment failure (TF) among population on antiretroviral therapy (ART) become a major public health threat, its magnitude of and factors leading to it are poorly defined. Hence, we aimed to estimate the magnitude of TF and its determinants in Ethiopia.

**Methods:** A follow-up study was conducted from March 2016 to 2017. Clinical and laboratory data were captured from paticipants’ medical record. Socio-demographics and explanatory variables were collected using structured questionnaire. Participants with baseline viral load (VL) >1000 copies/ml were followed for three to six month to clasify virologic failure (VF). Logistic regression was conducted to assess associated risk factors and statistical significance was set at P-value < 0.05.

**Results:** A total of 9,284 adults from 63 health facilities were included in the study.Viral load suppression (VLS) were found to be 8,180 (88.1%). Thirty-five percent of the study participants with VL>1000 copies/ml at baseline of the study were re-suppressed after three to six month of adherence counseling and hence VF was found to be 983 (11%). Immunologic and clinical failure was significantly improved from 21.5% and 16.5% at ART initiation to 576 (6.2%) and 470 (5.0%) at baseline of the study, respectively. Adherence, disclosure of HIV status, missed appointment to ART, history of ART exposure prior to initiation, residency and marital status had significant association with VLS.

**Conclusions:** VLS (88.1%) could explain the success of ART program in Ethiopia towards the UNAIDS global target. Eleven percent of the population is maintained on a failed first-line regimen. Improving adherence, ensuring disclosure of HIV status and appointment follow-up could improve treatment outcome.

## Introduction

For more than 35 years, the world has grappled with the AIDS pandemic that has claimed an estimated 35.0 million [28.9 million-41.5 million] lives and at its peak threatened global stability and security(1). Sub-Saharan Africa suffers from the undue burden of HIV, with an estimated 70% of the world HIV/AIDS infections and deaths occur in this region(2)(3). Ethiopia is one of the sub-Saharan African countries most affected by the HIV epidemic. According to the latest Spectrum modeling, an estimated 610,335 people were living with HIV in 2018(4).

Antiretroviral therapy (ART) is the lifesaving treatment for patients with HIV/AIDS, as it significantly reduces HIV-related mortality and morbidity(5).According to the report of the World Health Organization (WHO), the number of people receiving ART reached >20 million by mid-2017, up from about one million a decade ago(6). The impact of HIV infection has significantly decreased due to expansion of ART (7). The aim of the treatment is to suppress virus replication for as long as possible, restore and/or preserve immune function, improve quality of life, and reduce HIV-related morbidity and mortality(7). But this has not always been true due to treatment failure (TF), which can be explained by virological, clinical, or immunological causes.

Antiretroviral therapy has been consistently reported to suppress HIV-RNA to undetectable levels, and has decreased risk of clinical progression(2)(8). Despite these successes, increases in antiretroviral TF due to drug resistance and/or sub-optimal adherence to the regimen pose a major impediment to ART program. The WHO recommends viral load testing as the preferred approach to monitoring ART success and diagnosing treatment failure(9),(10).

WHO recommended that consecutive viral load measurements to confirm and define TF, among people receiving ART which should be performed 6 months after initiating ART and every 12 months thereafter(11), advirological failure (VF) is a more informative measure of TF(12). Monitoring individuals on ART for treatment failure remains the key challenges due to sub-optimal access to HIV viral load (VL) testing for routine follow-up of treatment (13,14). Current guidelines recommend that once patients start treatment, TF should not exceed 10%(11). However, studies show that TF is much higher than expected and viral replication continues to be a major challenge among patients living with HIV(15). Patients who experience TF will have increased risk of morbidity, mortality, on ward transmission as well as accumulation of drug resistance mutationswhen compared with patients with complete virological response(16). Previous studies revealed that various factors were associated with TF(7,16–18).

Appropriate response to TF requires data on population-level estimates of TF (virologic, immunologic and clinical outcome) to guide the national ART program to achieve successful treatment outcomes. Determining the magnitude of TF and identifying the associated risk factor of TF are of paramount importance to achieve a high treatment success rate and improve the quality of life of People living with HIV. However, there is limited evidence on the magnitude of TF and its determinants among HIV-infected patients on ART in Ethiopia. Thus, this study aimed to determine the magnitude of TF and identify associated risk factors among HIV-infected patients on ART in Ethiopia.

## Methods

### Study design and population

A retrospective and prospective follow-up study was conducted across the nationally representative health facilities (HFs) in Ethiopia. Baseline VL testing was done followed by second round VL testing after three to six months of intensive adherence counseling for patients with VL >1000 copies/ml at baseline to determine TF as defined by WHO-2013(10). Pre-established tools and questioner were also used to assess determinants of TF. HFs providing ART service for at least 60 patients and with at least 9 month of ART service experience was part of this study. Study participants within the HFs were those who had at least 9 month ART experience.

HFs to be included in this study was selected by using two stage cluster design. All HFs providing ART service in the country, by 2014, were listed (N=1,047). Following this, HFs serving for less than 60 ART patients was defined as facilities providing very small number of ART patients. These HFs were 415 in number and had for 7,992 (i.e. 2.47%) of the total ART patients in the country. Accordingly, 632 HFs have been included in this study.

According to the WHO recommendation, selection of 40 HFs is sufficient for nationally representative virologic failure (VF). However, due to the heterogeneity of the study participant load across the HFs and regions in Ethiopia, some regions and HFs with small number of ART patients may not be represented with this limited number of facilities to be sampled. Provided that one health center and one hospital should be represented across all the regional administrations in the country to maintain administrative balance, we included additional 23 HFs. Hence, the total numbers of HFs included in the study were 63.

Study participants were further stratified in to three based on the duration on ART: strata “A” (12±3 months), strata “B”(between 15 to 47 months) and strata “C” (at least 48 months).Moreover, it had also been reported that there is a 10% increment in TF across each respective stratum (19,20).

Magnitude of VF among strata “A” adult HIV-1 infected patients in seven university hospitals in Ethiopia is reported as 7% (unpublished data-EPHI). A confidence interval of half-width of ±5% has been used as appropriate compromise between feasibility and precision with 95% Confidence Interval (CI). Moreover, numbers of adults on ART within the HF across each stratum by the end of 2014 have been independently considered and calculated using the formula;

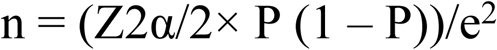

Where;

Z = value from standard normal distribution corresponding to desired confidence level (Z=1.96 for 95% CI)

P= Expected true proportion (0.07)

e= Desired precision (0.05)

Assumptions for sample size determination were: Proportion of patients on ART at different time point be 6% Strata “A”, 15% strata “B” and 79% strata “C” accordingly the population understudy would be 270,562; 317,443 and 343,511 respectively (21). Prevalence difference between each strata was also assumed to be 10% (19). Interview response rate and laboratory failure rate were considered to be 90% and15%(19) respectively. Considering 82.4% retention (22) and 10% non-respondent rate, a total sample size for adult population in this study is estimated to be 11,342.

### Variables of the study

Immunologic failure (IF), VF and clinical failure (CF) were considered to be dependent variables whereas socio-demographic variables, duration on ART, variable related to patient social behavior (disclosure, use of treatment assistant, use of memory aids, use of alternative medicine, use of alcohol and substance abuse, missed appointments), knowledge and perception on HIV and ART (knowledge and information on ART, Perception of treatment) and ART service delivery environment (waiting time, distance to the clinic, quality of care, trust in health care, providers, pill burden) were independent variables.

### Data collection

Data were collected in two rounds, to clasify the study population whether confirmed TF or not per the WHO classification (10).Base line data were captured for the study participants from March to August, 2016. Second-round data collectionwas specific for the study participants with baseline viral load >1000 copies/ml since December, 2016 to March,2017.

#### Types of data source

##### Primary data source

Pre-established questioner, used to capture data related with medication adherence, knowledge, attetude and perception on ART, service delivery enviroment and other demographic variables.

##### Secondary Data source

Seconday data were collected from the review of participants’ medical record. These variables included medical history at three points; by the time the participant start ART, the lettest record prior to the data collection and record during the data collection. Some of the key variables were CD4^+^ T-cell count, Clinical status per WHO clasification, Medication adherence history.

##### Blood sample collection

Ten milliliter blood sample was collected from the study participants during their attendance of the health facility for their appointment follow up. Plasma was separated after centrifugation and transported according to the standard operating procedure for sample collection, transport and tracking.

##### CD4^+^ T-cell count

CD4^+^ T-cell count was conducted at the health facility laboratory using eithr BD FACSCount ™ or BD FACS Calibour ™ (Becton Dickinson, USA).

##### Viral load testing

VL testing was conducted at base-line of the study and repeated for the study participants with baseline viral load >1000 copies/ml for the second round.

Viral load testing was conducted at the regional laboratories of the country using Abbott m2000sp system (Abbott Laboratories, Abbott Park, IL, USA) and COBAS® AmpliPrep/COBAS® Taq Man® HIV-1 Test, v2.0.

## Statistical analysis

The cumulative magnitude of VF was estimated from the proportion of adult with VL>1000 copies/ml both at baseline and second round of the study. Similarly, magnitude of immunologic and clinical failure were determined as per WHO definition(10).

Factors associated with VF was evaluated by comparing variables among adult who failed with those who never failed using the chi-square test for categorical data, and using student-t test for continuous variables. Logistic regression was done to determine the factors contributing to VF. The model were then built by dropping the most insignificant factor one at a time with factors whose P<0.05 were taken to be the factors that were independently associated with VF. All analyses were done using SPSS version 20.

### Operational definitions

- Adherence: The degree to which the person’s behavior corresponds with the agreed recommendations from a health care provider.
- ART Experience:

- Early stage ART experience: Population with ART experience from 9 to 15 months
- Mid-level ART experience: Population with ART experience from 16 to 47 months
- Advanced ART experience: Population with ART experience with at least 48 months
- Clinical Failure: New or recurrent clinical event indicating severe immunodeficiency (WHO clinical stage 4 condition) after 6 months of effective treatment
- Immunologic Failure:CD4 count falls to the baseline (or below) or persistent CD4 levels below 100 cells/mm^3^
- Virologic failure: Plasma viral load above 1000 copies/ ml based on two consecutive viral load measurements after 3 months, with adherence support
- Viral suppression: Plasma viral load above 1000 copies/ ml based on one viral load measurement

## Ethics statements

The project was ethically approved by EPHI Scientific and Ethics Review Office(SERO) before data collection. Confidentiality were respected during abstraction of data by the use of specific identification code for each enrolled patient number. Eligible study participants were identified by trained and experienced data collectors and supervisor at facility level. Information was provided about the study, and those willing to participate were provided with informed consent form to read and sign indicating that they understand the purpose of the study.

## Result

### Baseline characteristics

Of the 11,342 study participants, 11,013 (98.2%) responded to participate in the study, and 9,284 (84.3%) had successful VL result. Hence, the overall response rate at baseline of the study was 82.8%.

Table-1 shows demographic characteristics of HIV-infected people receiving ART. Demographically, 6,036(65.0%) were female and 3,234 (35.0%) were male. The median age of the study participants was 39 years, ranging from 15 to 90 years. Of the 9,284 participants,1,920 (21.0%) were under the age of 30 years and 2,351(25.3%) had no formal education and 3,937 (42.4%) attended elementary education. The mean ART experience of the study participants was 59 months, majority of them had at least 48 months ART experience that accounted for 5,559 (72.2%) followed by 16 to 47 months 1, 738 (22.6%) and 407 (5.3%).

**Table-1:**
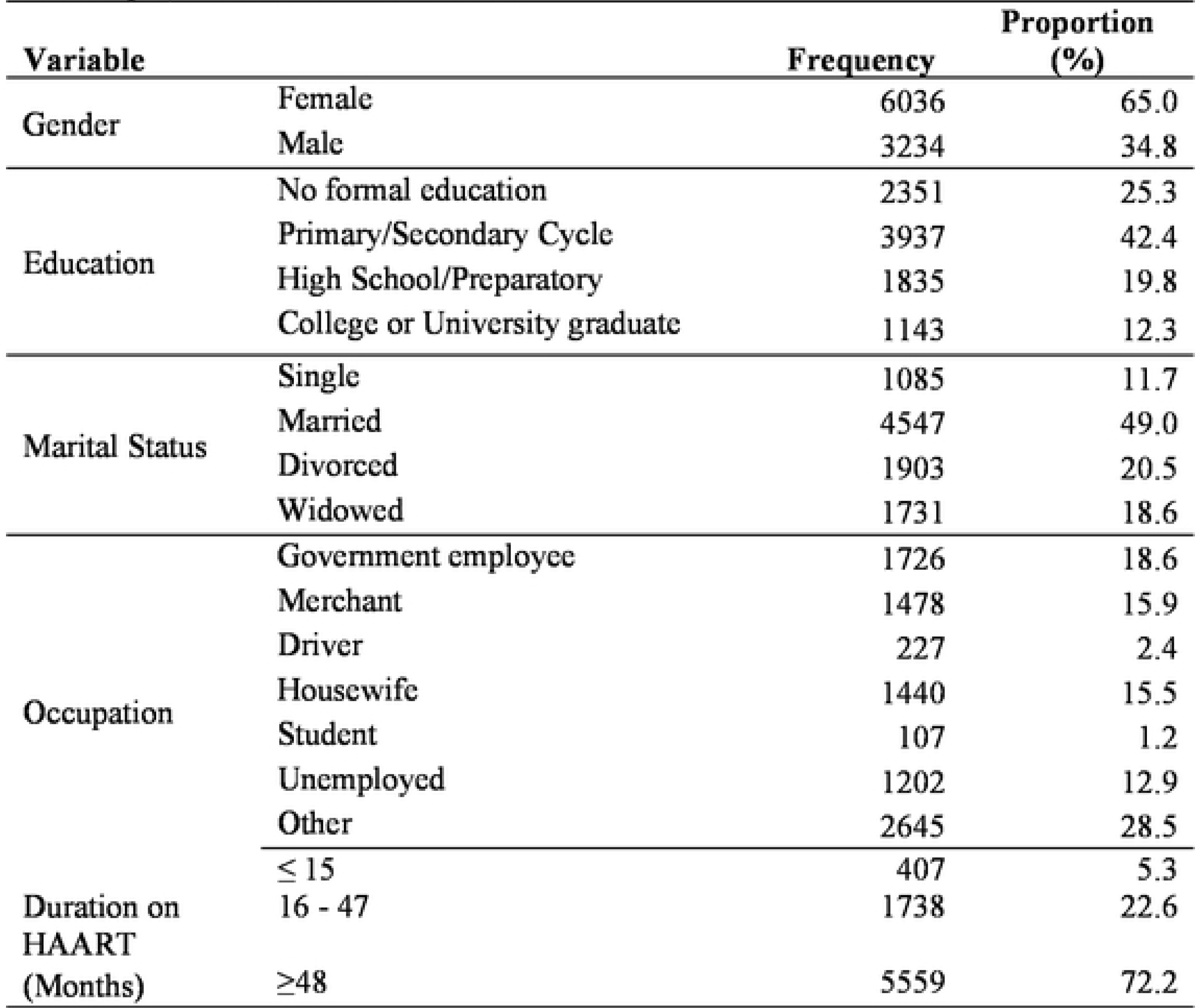
Socio-demographic characteristics ofHIV-infectedpopulation receiving ART in Ethioia, from March 2016 to 2017

| Variable |  | Frequency | Proportion (%) |
| --- | --- | --- | --- |
| Gender | Female | 6036 | 65.0 |
|  | Male | 3234 | 34.8 |
| Education | No formal education | 2351 | 25.3 |
|  | Primary/Secondary Cycle | 3937 | 42.4 |
|  | High School/Preparatory | 1835 | 19.8 |
|  | College or University graduate | 1143 | 12.3 |
| Marital Status | Single | 1085 | 11.7 |
|  | Married | 4547 | 49.0 |
|  | Divorced | 1903 | 20.5 |
|  | Widowed | 1731 | 18.6 |
| Occupation | Government employee | 1726 | 18.6 |
|  | Merchant | 1478 | 15.9 |
|  | Driver | 227 | 2.4 |
|  | Housewife | 1440 | 15.5 |
|  | Student | 107 | 1.2 |
|  | Unemployed | 1202 | 12.9 |
|  | Other | 2645 | 28.5 |
| Duration on HAART (Months) | ≤ 15 | 407 | 5.3 |
|  | 16 - 47 | 1738 | 22.6 |
|  | ≥48 | 5559 | 72.2 |

### Magnitude of treatment failure among population taking ART in Ethiopia

TF was classified according to WHO definition; VF is Plasma VL above 1000 copies/ ml based on two consecutive VL measurements after 3 to 6 months of the baseline, with adherence support. Immunologic failure was defined as CD4 count falls to the baseline (or below) or Persistent CD4 levels below 100 cells/mm^3^ and clinical failure is new or recurrent clinical event indicating severe immunodeficiency (WHO clinical stage 4 condition) after 6 months of effective treatment(10).

### Viral Load suppression among population taking ART in Ethiopia

The mean VL at baseline of the study was 1,190 copies/ml and the level of VLS among population taking ART in Ethiopia was 8,175 (88.1%) while1,109 (11.9%) had not achieved viral suppression. VLS at various level of ART experience: 9-15, 16-47 and ≥48 months were 90.20%, 87.85% and 84.2% respectively.

Overall, the proportion of viral suppression among the regions and city administrations in the country ranged from 81.5-90.3%. VLS significantly varied across regions in the country. The highest proportion of viral suppression was achieved in Southern Nations, Nationalities and people region (SNNPR) (90.3%) followed by Dire Dawa and Harari (89.6%, 89.4% respectively). On the other hand, participants from Somali region had the lowest viral suppression (81.5%) followed by Gambella (82.7%) and Addis Ababa (85.3%) (Table-2).

**Table-2:** Virological failure among HIV infected population receiving ART by region, from March 2016 to 2017

| Region | VLS | P-value |
| --- | --- | --- |
| Tigray | 88.56% | 0.00 |
| Dire Dawa | 89.55% | 0.61 |
| Addis Ababa | 85.29% | 0.95 |
| Afar | 93.50% | 0.02 |
| Amhara | 87.51% | 0.02 |
| Oromia | 88.30% | 0.28 |
| Somali | 81.52% | 0.51 |
| Benishangul Gumuz | 88.35% | 0.00 |
| SNNPR | 90.31% | 0.64 |
| Gambella | 82.72% | 0.57 |
| Hareri | 89.41% | 0.00 |

### Virologic Failure among population on ART in Ethiopia

Study participants with VL>1000 copies/ml at baseline of the study were recruited for the second round, with a dropout of 19%. Accordingly 895(81%) out of the 1,105 participants were included for the second round VL testing, done after 3 to 6 months of enhanced adherence counseling.Subsequently,316(35.3%) of the 895study participants were repressed during the second VL testing. Overall, the adjusted magnitude of VF among population taking ART in Ethiopia was found to be 11%.

### Immunologic and Clinical failure among population taking ART in Ethiopia

The mean CD4^+^ T-cell count among study participants during ART initiation and at baseline of the study improved from 193 to 482 cells/µl. The level of Immunologic failure among population taking ART in Ethiopia was found to be 6.2%, significantly improved compared to the time of ART initiation (21.5%). There was significant difference with regard to CD4^+^ T-cell count at ART initiation and at baseline of the study (P=0.001; 95% C.I. 3.21-4.50).

The level of clinical failure among population taking ART in Ethiopia improved from 16.5% at ART initiation to 0.5%.Clinical stage significantly improved at baseline of the study compared to the time when study participants were initiated for ART (P=0.003; 95% C.I. 2.91-3.24).

### Factors associated with viral suppression

Population taking ART living in rural Ethiopia was found to be at higher risk of VF with odds ratio (OR) 1.5 (95CI; 0.45-5.00; P=0.019). It was also found that, married individuals had higher VLS (95%CI; 0.31-1.24; OR=0.67; P=0.021).Individual factors; including, history of ART exposure before initiation, Alcohol use and disclosure of the HIV status had also association with viral suppression. Similarly, factors including, medication adherence, missed appointment and history of hospitalization also had association with viral suppression (Table-3).

**Table-3:**
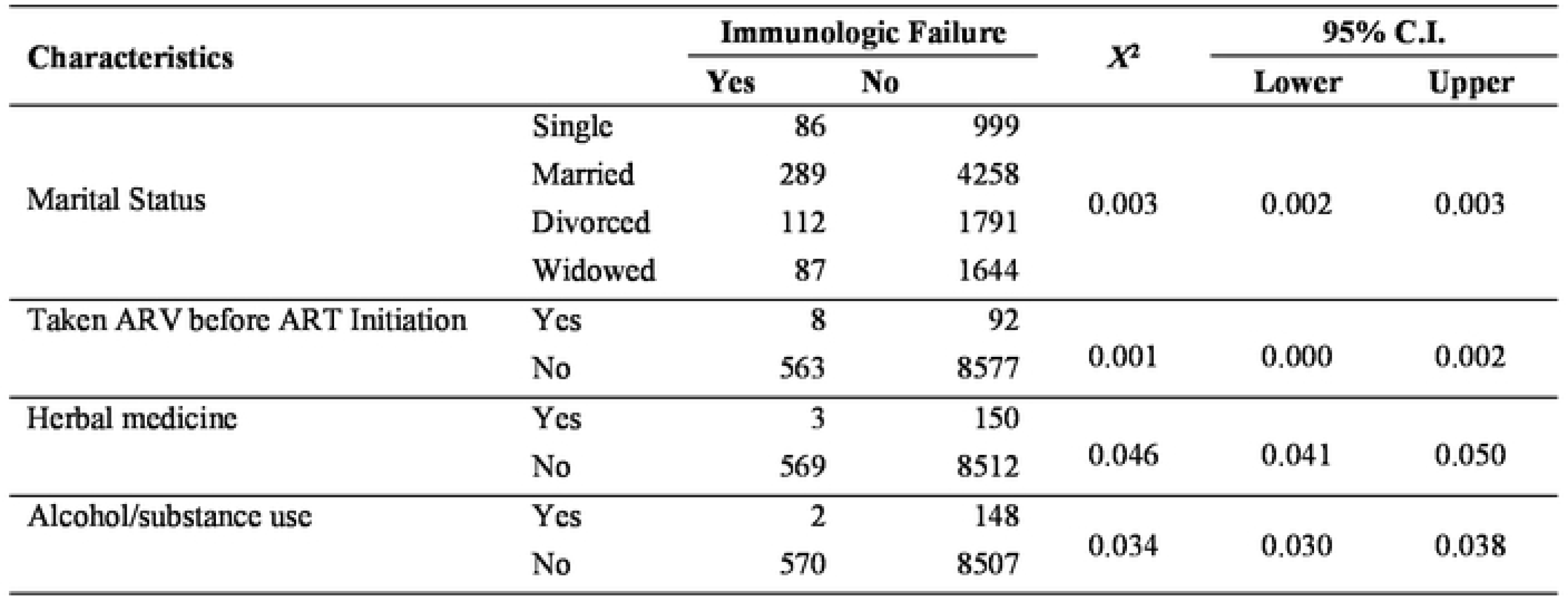
Socio demographic factors associated with virological failure among population taking ART in Ethiopia, from March 2016 to 2017

#### Factors associated with Immunologic failure

Marital status (p=0.003), previous history of ART exposure prior to ART initiation (P=0.001), history of use of herbal drugs (P=0.046) and alcohol use (P=0.034) had association with Immunologic failure (Table-5).

**Table-4:**
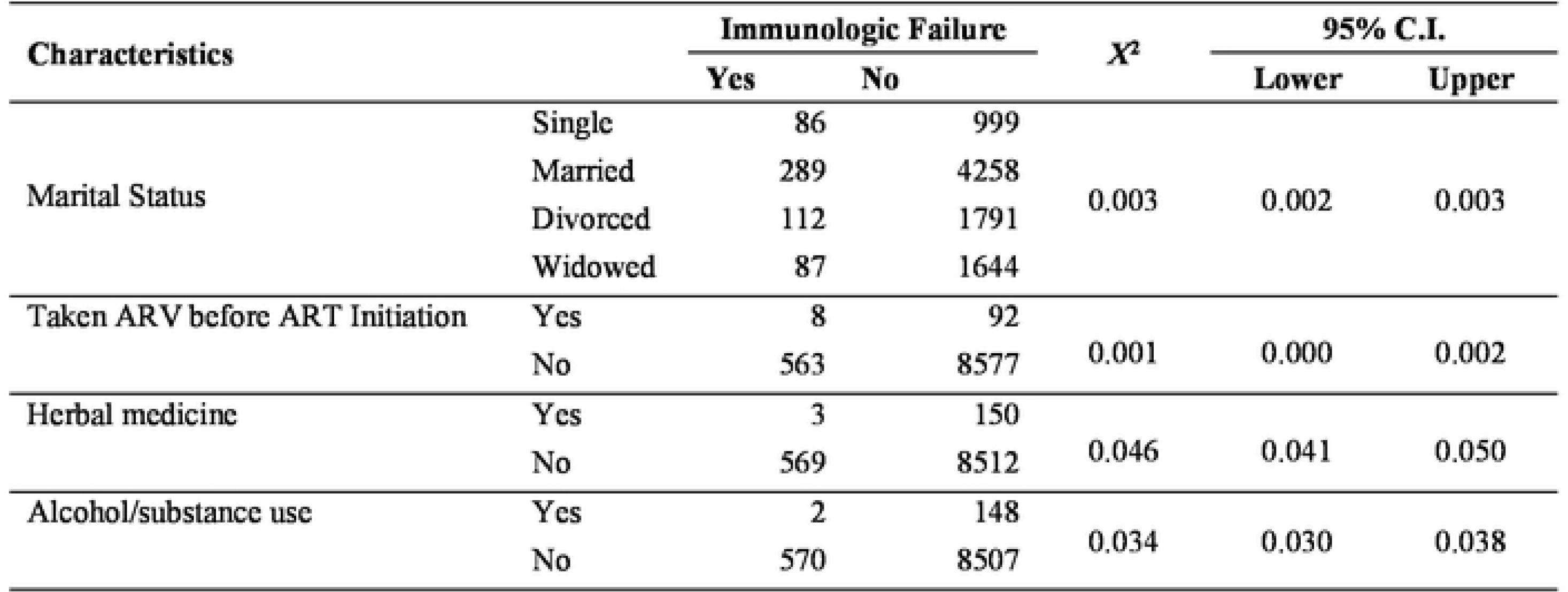
Factors Associated with IF among population on ART in Ethiopia, from March 2016 to 2017

**Table-5:**
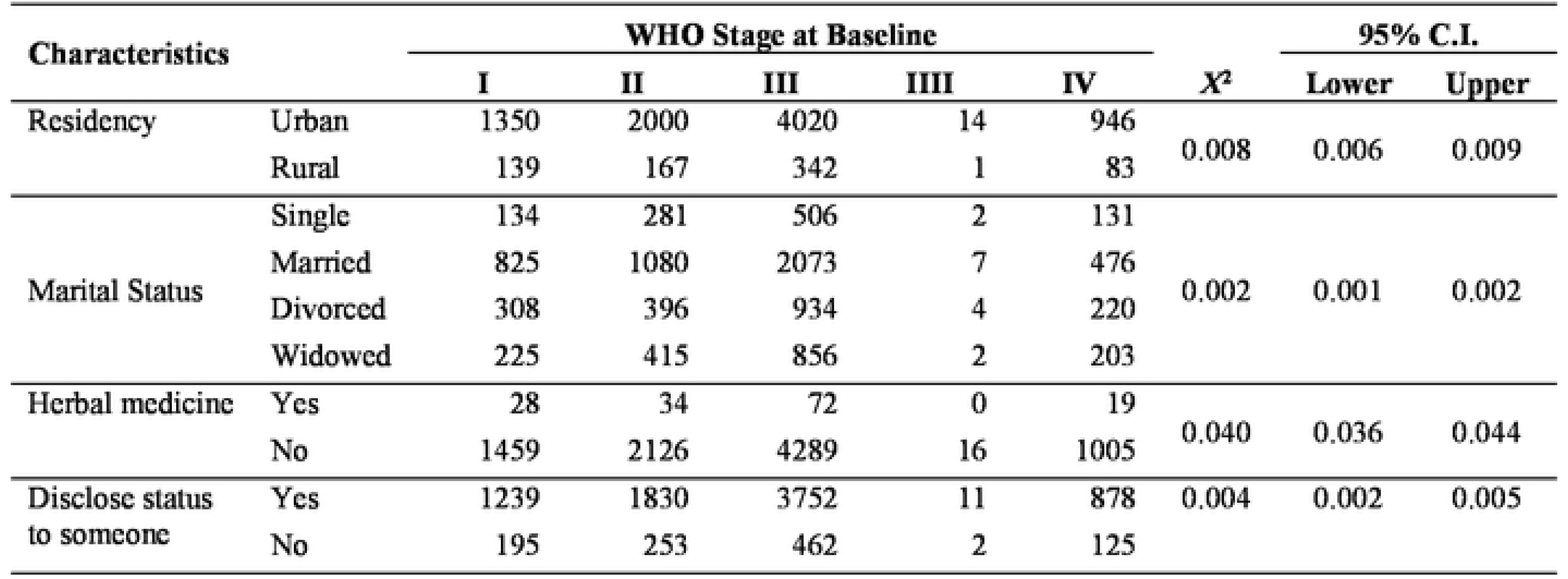
Factors associated with CF among population on ART in Ethiopia, from March 2016 to 2017

##### Factors associated with Clinical failure

Residency (P=0.008), Marital status (P=0.002), use of Herbal Medicine (P=0.040) and Disclosure status of HIV (P=0.004) had significant association with clinical failure (Table-5).

There were significant association between VF and IF (*X*^*2*^=3.11, P=0.001) and clinical failure had also signifiant association with VF (*X*^*2*^=6.446, P=0.03).

As limitation of the study, we could not verify whether the observed VF was due to resistance to ART drugs or not. Our data also showed considerable lost-to-follow up for the second round VL testing and missing data which might introduce some errors in our estimation.

## Discussion

In this nation-wide study, we evaluated the level of VF (using the definition of HIV-RNA VL>1000 copies/ml) amongst HIV infected adults receiving ART in Ethiopia and the determinants of VF. The findings of this study indicate the success of the ART program in the country. The finding of 11% VF in this study was lower than those studies from a number of lower than the figures from a number of low and middle income countries (10,17,23–26). The 88.1% VLS in this study is consistent with the finding of a previous cross-sectional study from Ethiopia in Tigray region which reported that TF was 11.5% while it is higher than a figure from a study conducted in Gondar(4.5%) (23,27). Despite having an overall high virological success, the level of virological suppression achieved in the subsequent viral load measurement among the patients with initial VF was low (35.3%). The possible explanation for such unfavorable prognosis after a long follow up could be due to the development of resistance to the current ARV drug and/or due to poor adherence to the treatment regimen. Our study supplemented the hypothesis in that, there was significant association between VLS and medication adherence.

This study also sought to compare regional variations in virological response of HIV patients. Our data showed a relatively higher VF among patients residing in Somali (18.5%), Gambella (17.3%), and Addis Ababa (14.7%). The least VF was noted among the patients from Afar region (6.5%). We have no clear explanation for such disparity in VF rate across the regions. Taking their geographical location into account, the observed VFs in Somali and Gambella regions might be partially explained by some logistic problems in getting pills on time as well as in obtaining psychological support. These might led to low adherence and the consequent VF. However, our speculation does not explain the least VF observed in Afar region, despite their similarity with the logistic issues, and to that of Addis Ababa too where the above mentioned challenges expected to contribute the least, if at all.

This study revealed a high level of HIV VLS among the study participants. About 88.1% of the study participants achieved virological suppression in the first round of the study. This is inline with the Ethiopian Population based HIV Impact Assessment (EPHIA) that reported 89%(10). This finding is also concurrent with the routine data from the country’s HIV program 87.3% (unpublished). VLS significantly varied by regional administrations in the country, which is also similar with the EPHIA report while there is limited disaggregated data at health facility level. This report is also in line with the estimation projection of the country’s 90-90-90 performance by UNAIDS (28). However, it is higher compared to a systematic review conducted in sub-Saharan Africa that reported VLS as 76% (29).

There was significant VL re-suppression after adherence and counseling for 3 to 6 months. Hence, confirmed VF among population taking ART in Ethiopia was found to be 11% which is the first report for the country. This finding contradicted with the report in HIV clinic of the coastal Kenya reported as 24.6%(17)and 6% (30) while it concurred with a study conducted in Uganda reported as 12.6% (31).

The finding from this study revealed that, the level of IF among population on ART in Ethiopia was 6.5% which is lower than that of a study conducted in Kenya which showed 13.3% while CF in this study indicated as5%which is in line with a similar study conducted in Kenya(5.7%)(30).

Most of the variables which found to have association with TF in this study were established facts to predict TF and are among the global recommendations to improve TF(8,32–34). Among these, medication adherence, disclosure of HIV status, missed appointment to ART, history of ART exposure prior to initiation, residency and marital had significant association with VLS which is in line with most of the studies conducted in low and middle income countries(32–36).

## Conclusion

TF among population taking ART in Ethiopia is still a public health concern, since 11% of virally failed population is maintained on failed first line regimen. The level of VLS (88.1%) could explain the programmatic success of Ethiopia towards achieving the UNAIDS global target of 90-90-90. Adherence to medication, disclosure of HIV status, regular follow-up to ART and care could significantly improve the treatment outcome of patients on ART in Ethiopia.

## Acknowledgements

We are grateful to the patients for participating in our study and to all health professionals and data collectors who involved in this work. We are very thankful to FHAPCO for providing technical support throughout the study.

